# Heat-Triggered Transcriptional Reprogramming in Microspores Disrupts Progression of Pollen Development in *Brassica napus* L

**DOI:** 10.1101/2024.11.15.623842

**Authors:** Neeta Lohani, Mohan B. Singh, Prem L. Bhalla

**Author notes:** **Correspondence:** Prem L Bhalla.

## Abstract

Rising temperatures due to climate change pose significant threats to global food security, particularly by impacting crop fertility and yield. Despite the known susceptibility of male reproductive development to heat stress, the specific adaptive mechanisms and critical regulatory genes involved in the stage-specific heat stress response in crops are still not well understood. This study examines the impact of acute heat stress on pollen development in Brassica napus (Canola), an essential oilseed crop, with a focus on the uninucleate microspore stage. We demonstrate that a brief exposure to 40°C during the uninucleate microspore stage results in loss of cellular polarity, failed asymmetric cell division, loss of cytoplasm, and pollen abortion, leading to a significant decline in pollen viability. Transcriptome analysis reveals the complexity of heat-induced transcriptional reprogramming in microspores, identifying 8.1% (6,245) of the expressed genes as responsive to high temperature, with a higher degree of downregulation. Fourteen differentially expressed transcription factors (TFs) belonging to diverse TF families were identified in heat-stressed microspores, with overrepresented target genes. The differential regulation of cell cycle control genes in heat-stressed microspores supports our microscopic observations and details the transcriptional reprogramming underlying the failure of heat-stressed microspores to undergo the first and second pollen mitotic divisions (PMI and PMII), resulting in aberrant pollen development in *B. napus*. Heat stress disrupts key biological pathways such as transcription, mRNA processing, translational control, protein homeostasis, autophagy, phytohormone signaling, and reactive oxygen species (ROS) homeostasis. Heat stress alters the intricate network of multiple signaling pathways, potentially disrupting the regulatory pathways underlying pollen development, with evident crosstalk underscoring the complexity of the heat stress response in pollen development. These findings underscore the critical vulnerability of the microspore stage to heat stress in Canola. Understanding the identified heat-responsive genes and regulatory networks provides valuable insights for breeding climate-resilient Canola varieties, thereby contributing to global food security amidst changing climatic conditions.

**Highlights:**

- Acute heat stress during microspore stage in *Brassica napus* disrupts pollen development.
- Heat stress induces complex transcriptional reprogramming in microspores, with higher degree of downregulation.
- Transcriptional control, RNA processing, coordination of protein homeostasis and translational control is altered in heat-stressed microspores
- Misregulation of cell cycle control genes supports the loss of cellular polarity and failure to undergo asymmetric cell division in heat stressed microspores.

## Introduction

Climate change has led to a substantial rise in global average temperature, with an increase of ∼1.1-1.2 since the late 19^th^ century. The decade from 2011 to 2020 set a new record as the hottest ever, with each of the past four decades surpassing the temperatures of all preceding decades since the year 1850 (NOAA, 2024). According to climate change projections, heatwaves will become more frequent and intense, and the affected land area is expected to quadruple by 2040 (Perkins-Kirkpatrick & Lewis, 2020). This presents a significant challenge to global food security, as high temperature events coinciding with critical developmental stages have a negative impact on crop productivity. The susceptibility of plants to high temperatures depends on the vulnerability of a specific developmental stage, which is species- and genotype-dependent (Janni et al., 2020; Prasad et al., 2017). However, the consensus regarding the heat stress sensitivity of crops identifies reproductive development especially male gametophyte development as most vulnerable to heat stress.

Male gametophyte development i.e., pollen development in flowering plants is complex and is divided into two stages: microsporogenesis and microgametogenesis (Borg et al., 2009; McCormick, 2004). Meiosis I and II occur during microsporogenesis, leading to the division of diploid pollen mother cells (meiocytes) to form four haploid uninucleate microspores. The microspores then undergo pollen mitosis I, producing two asymmetrically sized daughter cells - vegetative and generative cells - which have markedly different structural organization and developmental fates. The vegetative cell does not divide further and exits the cell cycle, whereas the generative cell undergoes symmetrical mitotic division (pollen mitosis II), resulting in the formation of two sperm cells. Depending on whether pollen mitosis II occurs before or after the pollen germinates, mature pollen is either bicellular or tricellular prior to release from the anther. Exposure to high temperature can impact different aspects of pollen development in crops. Heat stress can negatively affect recombination and cytokinesis during meiosis, disrupt tapetum development and function, result in the abortion of tetrads and developing microspores, lead to abnormalities in pollen morphology, reduce pollen viability and pollen germination rate thereby diminishing overall pollen function (Begcy et al., 2019; Fábián et al., 2024; Li et al., 2024; Lohani et al., 2022; Ning et al., 2021; Schindfessel et al., 2023; Q. Zhao, X. Guan, L. Zhou, M. A. U. Asad, et al., 2023). Thus, an in-depth understanding of the mechanisms underlying the male reproductive heat stress response in economically important crops is crucial to ensure future global food security.

*Brassica napus* L., commonly known as rapeseed, oilseed rape and canola, is one of the most important oil crops providing not only cooking oil for humans, but also protein-rich fodder for animals, and renewable materials for biodiesel and industrial applications. Canola is currently the second-highest yielding oil crop worldwide and accounted for 12.1% of global vegetable oil supply in 2021 (FAOSTAT, 2022). Canola seed yield losses ranging from 133-300 kg/ha is anticipated for every 1°C increase in the mean daily post anthesis temperature in Canada and Australia (Kutcher et al., 2010; Si & Walton, 2004). Dreccer et al. (2018) reported the association of higher maximum temperature after flowering with lower canola seed yield across major production zones in the 2009–2013 national variety trials in Australia. A meta-analysis of thirty-nine research studies, highlighted that early heat stress before end of flowering had the largest impact on canola seed yield and seed oil content (Secchi et al., 2023). Heat stress coinciding with reproductive stages in Canola can negatively impact the development of flowers, pollen fitness, fertilization as well as the synthesis and accumulation of oil, leading to compromised grain filling (Aksouh-Harradj et al., 2006; Angadi et al., 2000; Chen et al., 2021; Lohani et al., 2022; Pokharel et al., 2021; Young et al., 2004).

Several studies focussing on the impact of heat stress on Canola’s reproductive fitness followed various heat stress treatments ranging from 29-39°C lasting over a period of three to sixty days during different phases of reproductive development. For instance, Young et al. (2004) applied moderate heat stress of 35/18°C (day/night) with a gradual ramping up and down temperature cycle, lasting for one- or two-weeks during flowering, led to seedless or aborted pods. A two weeklong heat stress of 31/14°C (day/night), starting one week before the anthesis was reported to result in significant yield reductions in Canola (Koscielny et al., 2018). Pokharel et al. (2021) employed high day and high night temperature treatments (34/15°C, 23/20°C, 34/20°C) starting on the fourth day after first sign of flowering and lasting for two weeks and reported floral abnormalities and sterility leading to reduced seed yield and oil content. Although, such studies highlight the impact of different duration and intensity of high temperature exposure during different stages of reproductive development, on Canola’s seed production, the specific adaptive mechanisms to heat stress and the critical regulatory genes involved are still not well-understood in *B. napus*.

Crucial questions also remain concerning the underlying thermo-responsive molecular mechanisms during male reproductive development. Heat stress sensitivity and the impacts of transient heat stress during distinct pollen development stages have not been reported for *B. napus*. We recently investigated the effects of heat stress at 40°C for a period of 4h on mature pollen fitness and pollen-pistil interactions and revealed that acute heat stress impairs pollen viability, reduces pollen germination capabilities, and negatively impacts the ability of pistils to support pollen adhesion, germination and tube growth (Lohani et al., 2021). We further explored the changes in transcriptional patterns of mature pollen and pistil to unravel the molecular signatures in response to acute heat stress (40°C for 30 min). High temperature altered key regulatory genes involved in unfolded protein response, transcriptional networks, pollen-pistil interactions, cellular organization, transport, and metabolism leading to disruption of successful fertilization.

To further explore the impact of acute heat stress on male reproductive development in a stage-specific manner, we investigated the effect of a single episode of extreme heat stress during uninucleate microspore stage on progression of pollen development at the morphological level. We further performed an *in silico* transcriptomic analysis to capture the heat stress triggered transcriptional reprogramming in uninucleate microspores resulting in disruption of pollen development and pollen abortion. We explored the heat stress responsive pathways, such as heat stress signalling, phytohormone regulation, cell-cycle progression, and the transcriptional and translational machinery in uninucleate microspores and highlighted that different stages of male reproductive development have specific heat stress responsive molecular signatures. This dataset provides the foundation for the discovery of essential regulatory pathways and genes for imparting developmental stage specific heat tolerance, leading to the development of climate-smart Canola varieties.

## Materials and Methods

### Plant growth conditions and temperature treatments

A commercial cultivar of *B. napus* (AV Garnet) was selected for this study. The plants were grown and maintained in Thermoline growth cabinets located within the Plant Growth Facility at the University of Melbourne, Australia under controlled growth conditions of 23/18 temperature, 200 μmolm^-2^s^-1^ light intensity (16h light/8h dark) and 60% humidity. Around 50-55 days after sowing (das) the plants bearing secondary inflorescences were subjected to heat stress treatment at 40°C (single exposure) for four hours (0900 to 1300h). Prior to heat stress treatment, buds > 1.5mm were manually removed from at least five secondary inflorescences per plant (**Figure S1A**) to only keep the buds predominantly at uninucleate microspores stage in the outermost whorl of the inflorescence and these inflorescences were tagged. The plants were well-watered to prevent the onset of drought stress, soluble nutrient fertilizer was applied directly to the soil once a week for each plant and the plants were spatially randomized weekly. The experiments were conducted with three biological replicates, with each replicate consisting of twenty-five plants.

### Sample preparation and microscopy

DAPI staining was performed to detect the DNA and track the progression of pollen development from uninucleate microspores to mature tricellular pollen. The samples were observed under a fluorescence microscope (Olympus BX60) using the appropriate filter depending on the staining method.

Pollen viability was assessed through a double staining technique using Fluorescein Diacetate (FDA) and Propidium Iodide (PI), as previously described by Lohani et al. (2022). Briefly, mature pollen grains were released from anthers of appropriate size (6–7 mm) by gently macerating the anthers in a staining solution containing 10% sucrose, 2 mgmL^-1^ FDA, and 1 mgmL^-1^ PI. The samples were then incubated in the dark for 20 min at room temperature and observed under a fluorescence microscope (Olympus BX60). Acetocarmine staining was performed to differentiate between viable pollen and aborted pollen.

For cross sections, microspore containing buds exposed to 40 for 4h were collected after five days of heat stress treatment at the mature bud stage (6-7 mm). The buds were fixed overnight in Carnoy’s solution and then in 70% ethanol for long term storage until they were processed for paraffin embedding. Embedded buds were cut into 5µm thin cross sections using a vibrating blade microtome (Leica VT1000 S). Paraffin sections were dried overnight, dewaxed, and rehydrated before staining with toluidine blue for morphological observations.

### Isolation of uninucleate microspores

Flower buds (1-1.5 mm) containing uninucleate microspores were collected from non-stressed plants grown under optimal growth conditions and from heat stressed plants treated at 40°C for 30 minutes. The buds were collected immediately at the end of the single-exposure heat stress treatment. The buds were placed in pre-chilled 0.5×B5 medium in a Petri dish kept on ice. The anthers were carefully dissected from the buds and then macerated in the B5 medium. This crushed suspension was then filtered using a 44µm nylon mesh into 1.5 mL Eppendorf tubes and the filtrate was centrifuged for three minutes at 150 g at 4°C. The supernatant was discarded, the pellet was washed with 0.5×B5 medium, and the centrifugation process was repeated. This entire step was repeated twice. Following removal of the supernatant, the pellets immediately frozen in liquid nitrogen and stored at -80°C. After the isolation of uninucleate microspores, an aliquot from the isolated pellet was used to perform DAPI staining to check the purity of the samples. The samples were 85% enriched in uninucleate microspores. The remaining 10-15% of the sample consisted of tapetum cells, anther wall components and other non-microspore debris.

### RNA extraction, library preparation and RNA-Sequencing

Total RNA was isolated from both non-stressed and heat-stressed microspore samples was using mirVana™ microRNA isolation kit in accordance with the manufacturer’s instructions. These RNA samples were then stored in dry ice and shipped to BGI Tech Solutions (Hongkong) CO. Limited, where they were subjected to quality control protocols for RNA sequencing. RNA sample integrity was tested using an Agilent Bioanalyzer 2100™; this confirmed their suitability for being sequenced using the BGISEQ-500 platform for PE100 strand-specific mRNA sequencing; with a sequencing depth of 30 million raw reads for each sample. Following sequencing, data filtration was performed, which included removal of adaptor sequences, contamination, and poor-quality reads.

### Differential expression analysis

FastQC v0.11.8 (Andrews, 2010) was employed to perform quality checks on the raw sequencing files. Reference transcriptome file for *B. napus* was downloaded from Genoscope (http://www.genoscope.cns.fr/brassicanapus/data/). Transcript expression was quantified using Kallisto v0.44.0 (Bray et al., 2016). Conversion of transcript expression levels to gene expression levels was carried out using tximport v1.6.0 (countsFromAbundance=“no”) (Soneson et al., 2015). Pre-filtering of low-count genes was undertaken by retaining the genes with a minimum of five total counts across the samples. Differential expression analysis was performed using DESeq2 R package v1.28.1. DESeq2 was loaded with tximport data using DESeqDataSetFromTximport, which created an offset and corrected for alterations to the average transcript length across the samples (Love et al., 2014). Principal component analysis (PCA) was performed to determine the relatedness of biological replicates. For the generation of log_2_ foldchange estimates with greater accuracy, the lfcShrink (type = “apeglm”) function was employed. The differential expression thresholds were established as fold change 1.5 and p-adjusted value cut off 0.01 (lfcthreshold = 0.585, altHypothesis = “greaterAbs”, alpha = 0.01, pAdjustMethod=“BH”) for the alternate hypothesis, BH: Benjamini-Hochberg (Love et al., 2014; Zhu et al., 2019).

### Functional analysis and visualisation

The differentially expressed genes were functionally annotated by combining the outputs of using the PlantRegMap Platform (Tian et al., 2020) and PANNZER2 tool (Törönen et al., 2018). The GO enrichment was performed using *topGO* R package (Alexa & Rahnenführer, 2009). Subsequently, pathway annotation and enrichment analysis of differentially expressed genes was carried out using the KOBAS 3.0 database (Bu et al., 2021). A GO term and a KEGG pathway were considered significantly enriched only when the corrected p-value for that pathway was <0.01 after applying Fisher’s exact test and false discovery rate (FDR; BH method) correction. Furthermore, homologous *B. napus* genes, in comparison with the Arabidopsis proteome, were identified using the BlastP program with an E-value ≤ 1e-05. We also performed an orthology analysis between *B. napus* and *Arabidopsis thaliana* genes using *g:ortho* a function of *g:profiler* (Raudvere et al., 2019). We transferred the gene descriptors annotated by PANNZER2 tool to differentially expressed genes. To identify Transcription Factor Binding Sites (TFBS) in the differentially expressed gene sets we employed SeqEnrich (Becker et al., 2017). Visualization of the significantly enriched functional pathways was performed using the *ggplot2* R package (Wickham, 2011) and plotting tools in MATLAB R2021a. R packed *ComplexHeatmap* package was used to generate gene expression heat maps (Gu & Hübschmann, 2022). Gene regulatory network visualization was performed using Cytoscape 3.10.1. Transcripts per kilobase million (TPM) values were used for normalisation of gene-specific read counts and Z-score values i.e., scaled TPM values were used to represent gene abundance in heatmaps.

## Results and Discussion

### High Temperatures during Microspore Stage Results in Aberrant Pollen Development

To investigate the effects of high temperature on the pollen development program in *B. napus,* we subjected the plants to a single four-hour episode of heat stress at 40°C, followed by returning the plants to optimal growth temperatures (23/18°C light/dark). A non-stressed/control group of *B. napus* plants was maintained under the optimal conditions for comparison. Prior to heat stress treatment, >1.5 mm buds were removed from secondary inflorescences and these inflorescences were tagged in both sets of plants (**Figure S1A**). The buds in the outermost whorl of tagged inflorescences were predominantly at the uninucleate microspore stage of pollen development. Following heat stress treatment, we tracked the progression of microgametogenesis in both non-stressed and heat-stressed plants. Microscopic examination revealed that pollen development proceeded normally in non-stressed plants (**Figure 1A**), where uninucleate microspores underwent polarization and pollen mitosis I (PMI, asymmetric division), resulting in bicellular pollen with two unequal daughter cells: a larger vegetative cell and a smaller generative cell. The generative cell proceeded through pollen mitosis II (PMII) to form two sperm cells resulting in mature tricellular pollen. Conversely, heat stress had a pronounced negative impact on uninucleate microspores, leading to abnormal progression of pollen development. Three days after heat stress, the stressed microspores showed loss of cellular polarity followed by unsuccessful asymmetric mitotic division. As a result of heat stress induced aberrations, the mitotic division led to generative and vegetative cell of similar sizes. The vegetative cell failed to undergo PMII leading to pollen collapse, loss of cytoplasm and pollen abortion by the seventh day (**Figure 1B**).

**Figure 1.**
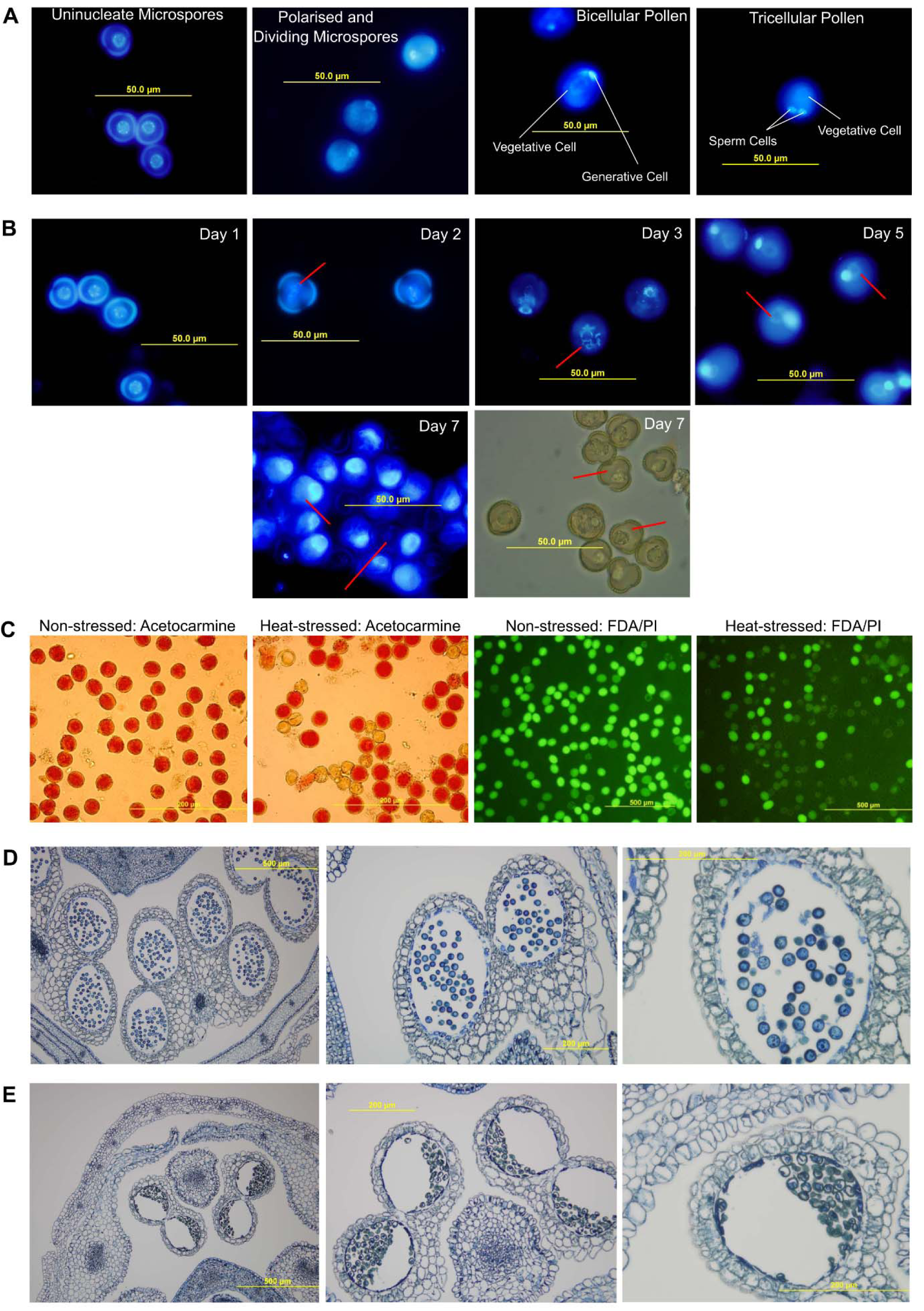
**(A)** Representative images of DAPI staining illustrating the progression of pollen development from the uninucleate microspore stage to the mature tricellular pollen under non-stressed conditions in *B. napus*. Uninucleate microspores undergo polarization and then form bicellular pollen by pollen mitosis I which results in two unequal daughter cells, the bigger vegetative cell, and a smaller generative cell. The generative cell further undergoes pollen mitosis II to form two sperm cells leading to the formation of a tricellular mature pollen. **(B)** Representative images of pollen development stages in buds heat-stressed during the uninucleate microspore stage at 40°C for four hours. Staining performed on Day 1, Day 2, Day 3, Day 5, and Day 7 after heat stress treatment. In the last panel, bright field image taken on Day 7 after heat stress treatment highlight the absence of cytoplasm and collapsed structure of the microspores which failed to develop into mature tricellular pollen. The red lines are highlighting the irregularities in progression of pollen development. **(C)** Pollen viability test by acetocarmine staining and FDA/PI staining of pollen isolated from mature buds from non-stressed and heat-stressed plants. **(D)** Cross sections of 6-7 mm *B. napus* buds. Representative aniline blue stained cross section images of anthers from non-stressed buds as viewed via bright field. In non-stressed bud sections, normally developed pollen was observed in the anther locules. **(E)** Cross sections of 6-7mm *B. napus* buds. Representative aniline blue stained cross section images of anthers from bud’s heat-stressed during uninucleate microspore stage, as viewed via bright field. Due to the heat stress exposure during uninucleate stage, there are some irregularities in tapetal degradation, the developed tricellular pollen was aborted and was collapsed and shrunken.

Microscopic observations highlighted the negative effect of short-term heat stress on pollen development, prompting further investigation of the effects of heat stress during the uninucleate microspore stage on pollen fertility. A significant decline of 60-65% in pollen fertility was observed in mature buds collected from heat-stressed plants, indicating failure of nearly two-thirds of the heat-stressed microspores in fulfilling their developmental fate to form tricellular pollen (**Figure 1C**). Apart from a few studies, the effects of heat stress on progression of pollen development from uninucleate microspores to tri-cellular pollen have not been systematically described in plants. In maize, heat stress treatment at 35/25°C (light/dark) for a period of 48h during the unicellular stage resulted in accelerated pollen development and reduced pollen germination rate (Li et al., 2024). Similarly, exposure to heat stress at 39/30°C (day/night) for two to four days during early microspore stage in rice results in a decline in pollen viability and germination (Endo et al., 2009). Cross sections of non-stressed (**Figure 1D**) and heat-stressed (**Figure 1E**) anthers in *B. napus* also revealed heat stress induced pollen abortion and abnormal degradation of tapetal cells in the anther locules. Notably, pollen developmental defects have been attributed to delayed tapetum degradation through programmed cell death in several plants (Djanaguiraman et al., 2018; Fábián et al., 2024; Oshino et al., 2007; Suzuki et al., 2001; Q. Zhao, X. Guan, L. Zhou, M. A. U. Asad, et al., 2023). Collectively, these findings highlight that even short-term heat stress exposure during a critical stage of male reproductive development can severely disrupt the developmental program and significantly reduce male fertility in *B. napus*.

### Heat Stress Induces Complex Transcriptional Reprogramming in Microspores

To investigate the molecular basis underlying heat induced abnormal male reproductive development in *B. napus*, we employed a strand-specific RNA-sequencing approach to compare the transcriptome of heat-stressed and non-stressed uninucleate microspores. The non-stressed and heat-stressed samples showed significant biological relatedness (**Figure S1B**) with an average pairwise correlation coefficient of 0.92 and 0.98, respectively. Differential gene expression analysis revealed that 8.1% (6,245; 2604 upregulated and 3641 downregulated) of the expressed genes (77,147) were responsive to high temperature in microspores (**Figure 2A**, **Table S1**). In our previously published study, only ∼3% of expressed *B. napus* genes (47,779) were identified as differentially regulated in heat-stressed mature pollen (Lohani et al., 2021). The higher percentage of expressed genes and heat stress responsive differentially regulated genes in microspores compared to mature pollen is expected as decreasing trend in transcriptome size and complexity during male germline development is extensively reported in flowering plants (Cui et al., 2015; Rutley & Twell, 2015; Wei et al., 2010). Moreover, in heat-stressed microspores, a greater percentage of genes were downregulated in response to high temperature compared to mature pollen. This pattern of differential expression in the heat-responsive transcriptome between post-meiotic pollen and mature pollen has also been reported in tomatoes (Keller & Simm, 2018).

**Figure 2.**
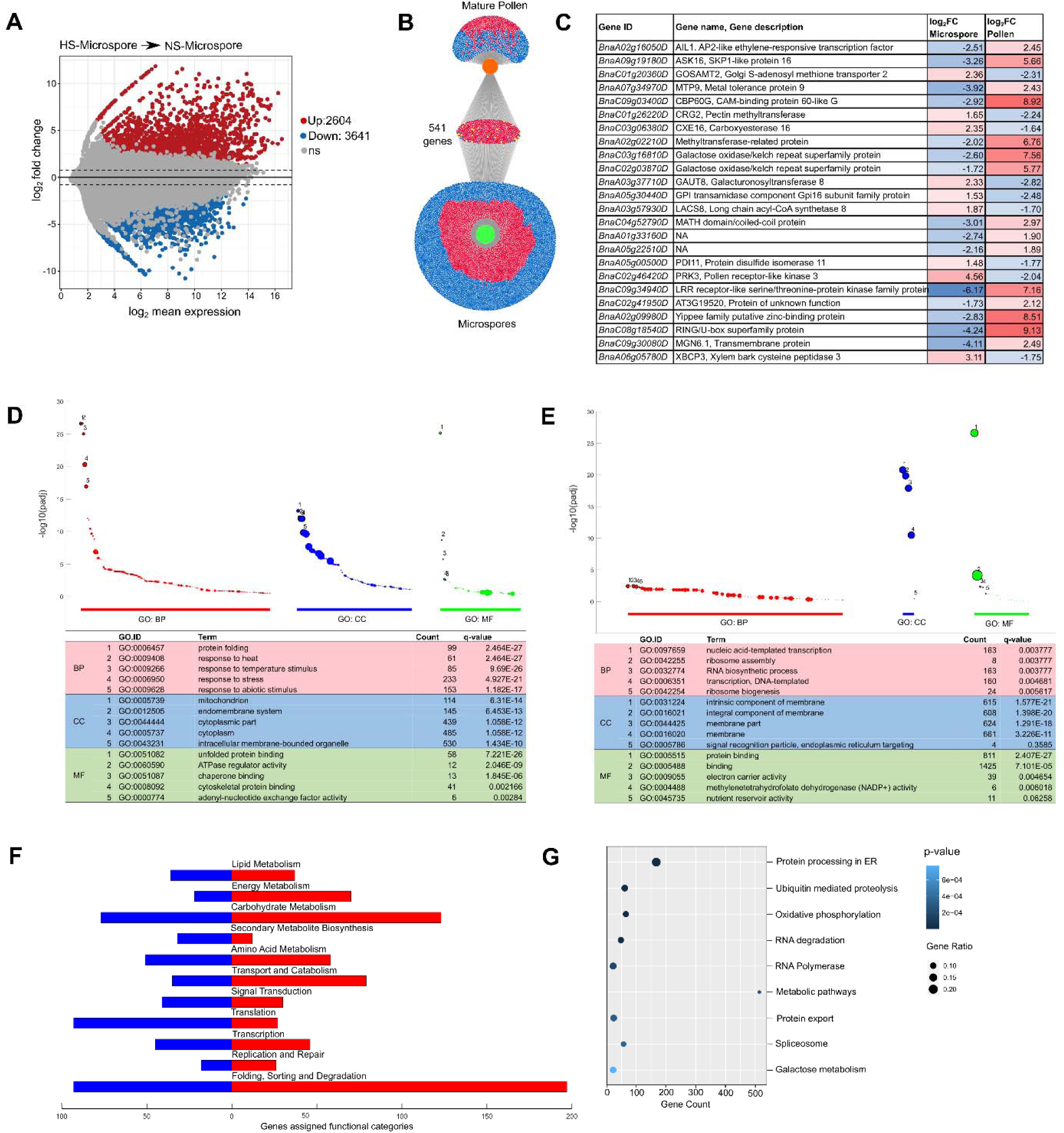
**(A)** MA Plot of the differentially expressed genes in heat-stressed microspores; red dots represent upregulated genes; blue dot represents downregulated genes; and grey dots represent non-significant genes. **(B)** Diagram representing the intersecting and specific differential regulation of heat responsive transcriptome between microspore and mature pollen. Blue dots: downregulated, red dots: upregulated, yellow dots: up- or down-regulated. **(C)** Detailed summary of the 24 heat responsive genes regulated in opposite manner in mature pollen and microspores in response to heat stress. GO term enrichment analysis of upregulated **(E)** and downregulated **(F)** differentially expressed genes in heat-stressed microspore. GO terms belonging to the three categories Biological Processes (BP), Cellular Component (CC) and Molecular Function (MF); are represented in the figure. The dot size is proportional to the number of genes included in the term, and the top five significant terms are highlighted. **(G)** KEGG Pathway annotation of differentially expressed genes in heat-stressed microspores. Distribution of genes in each functional category is represented by red if upregulated and blue if downregulated in response to heat stress. **(H)** Enriched KEGG pathways (p < 0.01) associated with the differentially expressed genes in heat-stressed microspores. HS: heat-stressed, NS: non-stressed.

Further comparative analysis of the transcriptomes of mature pollen and microspores revealed an overlap of 541 heat-responsive genes (**Figure 2B**, **S1C**). Of these, 517 genes exhibited similar patterns of differential regulation, with 500 upregulated and 17 downregulated. Conversely, 24 genes displayed contrasting regulation patterns (**Figure 2B**, **2C**). Protein similarity analysis of these 24 genes against Arabidopsis showed that the majority have not been functionally validated for their potential roles in heat stress response and/or pollen development. Among these 24 genes, *BnaA05g00500D*, identified as a homolog of the ARABIDOPSIS PROTEIN DISULFIDE ISOMERASE 11 (PDI11) gene, is an endoplasmic reticulum (ER)-localized protein and is upregulated in response to ER stress (Fan et al., 2021; Yuen et al., 2013). ER stress plays a crucial role not only in the heat stress response but also in male gametophyte development in plants (Singh et al., 2021). The differential expression patterns of *BnaA05g00500D* between mature pollen and microspores under heat stress suggest that this gene may regulate both pollen development and the heat stress response in a stage-specific manner. PDIs are key players in maintaining ER homeostasis and ensuring correct protein folding, which are essential for protein stability, enzymatic function, cellular transport and unfolded protein response (UPR) (Bhave et al., 2011; Laurindo et al., 2008; Lu & Christopher, 2008). In Arabidopsis, the ER-localized PDI9 protein plays a key role in controlling both pollen development and its tolerance to high temperatures (Feldeverd et al., 2020). Similarly, in rice, several PDI genes are upregulated in heat-stressed anthers, and silencing of the OsPDIL1-1 gene increased NADPH oxidase activity, leading to the excessive production of reactive oxygen species (ROS). This overaccumulation of ROS negatively impacted pollen viability and made floret fertility more vulnerable to heat stress (Q. Zhao, X. Guan, L. Zhou, Y. Xu, et al., 2023). Together, these findings suggest that PDIs, including *BnaA05g00500D*, may have a conserved role across species in mediating the stress response and reproductive development under heat stress conditions, particularly by modulating pollen viability and function. Further research is needed to investigate the functions of these genes and to elucidate the molecular mechanisms underlying the transcriptional machinery related to development and heat stress response at different stages of pollen development.

The differentially expressed genes in heat-stressed microspores were annotated by integrating multiple approaches: BLASTP analysis against the Arabidopsis proteome (threshold of e-value 1e-5), orthology analysis between Arabidopsis and *B. napus* proteins, functional annotation using PANNZER2, Gene Ontology (GO) term and KEGG pathway analysis (**Table S2**). Several genes remained unannotated, indicating genome expansion in the *Brassicaceae* family over the course of evolution. Among the top 20 up-regulated genes, homologs of gene encoding for Heat Shock Proteins (HSP15, HSP17, HSP21, HSP23, HSP90) and DnaJ Chaperone Protein were identified as key heat stress responsive genes. Additionally, a homolog of BETA-GALACTOSIDASE 15 (BGAL15) was significantly up-regulated in heat-stressed microspores. BGAL, known for its role in pollen development, acts on various substrates, including arabinogalactans, galactolipids, and pectin, to release galactose, aiding cell wall loosening after microspore mitosis during pollen development (Hruba et al., 2005). Analysis of the top 20 down regulated genes revealed genes potentially encoding for EMBRYO DEFECTIVE 3127 (EMB3127), LATE EMBRYOGENESIS ABUNDANT 3 (LEA3), Histone H2B.2, and GIBBERELLIN 20 OXIDASE 5 (GA20OX5). In Arabidopsis, EMB3127 is reported to a play role in cell cycle control (Meinke, 2020), LEA3 is identified as a stress responsive gene (Luhua et al., 2013), Histone H2B.2 is involved in chromatin remodelling (Zarreen et al., 2022) and GA20OX5 is associated with gibberellic acid signalling (Lee et al., 2016).

Furthermore, enrichment analysis of GO terms (**Figure 2D**, **2E**) and KEGG pathways (**Figure 2F**, **2G**) identified genes associated with response to stress, protein homeostasis, transcription, RNA processing, translation, signal transduction and metabolism. Thus, differential gene expression analysis mapped out a complex network of transcriptome reprogramming in microspores triggered by heat stress.

### Acute Heat Stress at the Microspore Stage Induced Differential Regulation of Transcription Factors and RNA Processing Machinery

Gene expression patterns are strongly regulated by transcription factors (TFs). Deviations from baseline conditions, resulting in significant transcriptional modifications are generally attributed to alterations in TF activity. We performed a TF prediction and identified 5.3% of the differentially expressed genes i.e., 328 genes as TFs belonging to 43 TF gene families (**Table S3A**). This indicates a wide-ranging and complex disruption in the transcriptional regulation in microspores exposed to heat stress. Specifically, we found 157 TFs were upregulated, and 171 were downregulated, with key stress-responsive TF families such as bZIP, AP2, ARF, BES1, BBR-BPC, DOF, MYB, NAC, WRKY, ERF, GRAS, and others showing differential regulation (**Figure S2**). Additionally, the number of TFs expressed in microspores is significantly higher than mature pollen with only 23 heat stress responsive TFs belonging to 18 TF families (Lohani et al., 2021). This difference is obvious as the transcriptional machinery is much more complex during microspore stage of male gametophyte development.

We performed a TF enrichment analysis and identified 14 differentially expressed TFs with overrepresented target genes across the differentially expressed gene set in heat-stressed microspores. Homologs of HEAT STRESS FACTOR B2A (HSFB2A), HSFB2B, bZIP28, G-BOX BINDING FACTOR 3 (GBF3), NAC DOMAIN CONTAINING PROTEIN 17 (NAC017), NAC002, BASIC HELIX-LOOP-HELIX 80 (bHLH80), MYB DOMAIN PROTEIN 77 (MYB77), TARGET OF EARLY ACTIVATION TAGGED 2 (TOE2) and REVEILLE 7 (RVE7) were upregulated, whereas HSFA4A, bZIP60, NAC069 and TCP DOMAIN PROTEIN 2 (TCP2) were downregulated in heat-stressed microspores (**Table S3B**). We further identified the biological processes associated with their target genes and built an enriched gene regulatory network which is activated by high temperature in microspores (**Figure 3A**). The network highlights common and specific biological processes potentially regulated by these TFs. HSFs are identified as central regulators of heat stress response. HSFB2A, and HSFB2B regulate the expression of other HSFs and HSPs and are also required for reproductive development and thermotolerance (Wunderlich et al., 2014; Zhang et al., 2024). In heat-stressed microspores, the target genes of HSFB2A and HSFB2B commonly regulated genes associated with response to stress, protein folding and response to oxidative stress among other biological processes (**Figure 3A**). Interestingly, all the HSFs were upregulated in heat-stressed microspores, except for HSFA4A. The target genes of HSFA4A were not only associated with genes involved in response to stress but also pollen development. However, HSFA4A has not been functionally validated to be associated with reproductive development in plants. In *B. napus*, *BnHSFA4A* regulates the activity and expression of GALACTINOL SYNTHASE (GOLS) genes and thereby triggers an antioxidant system including other stress responsive mechanisms in *B. napus* (Lang et al., 2017). Interestingly, GOLS1 was significantly up-regulated in heat stressed microspores highlighting the role of a HSFA4A-GOLS1 regulatory module in male reproductive heat stress response.

**Figure 3.**
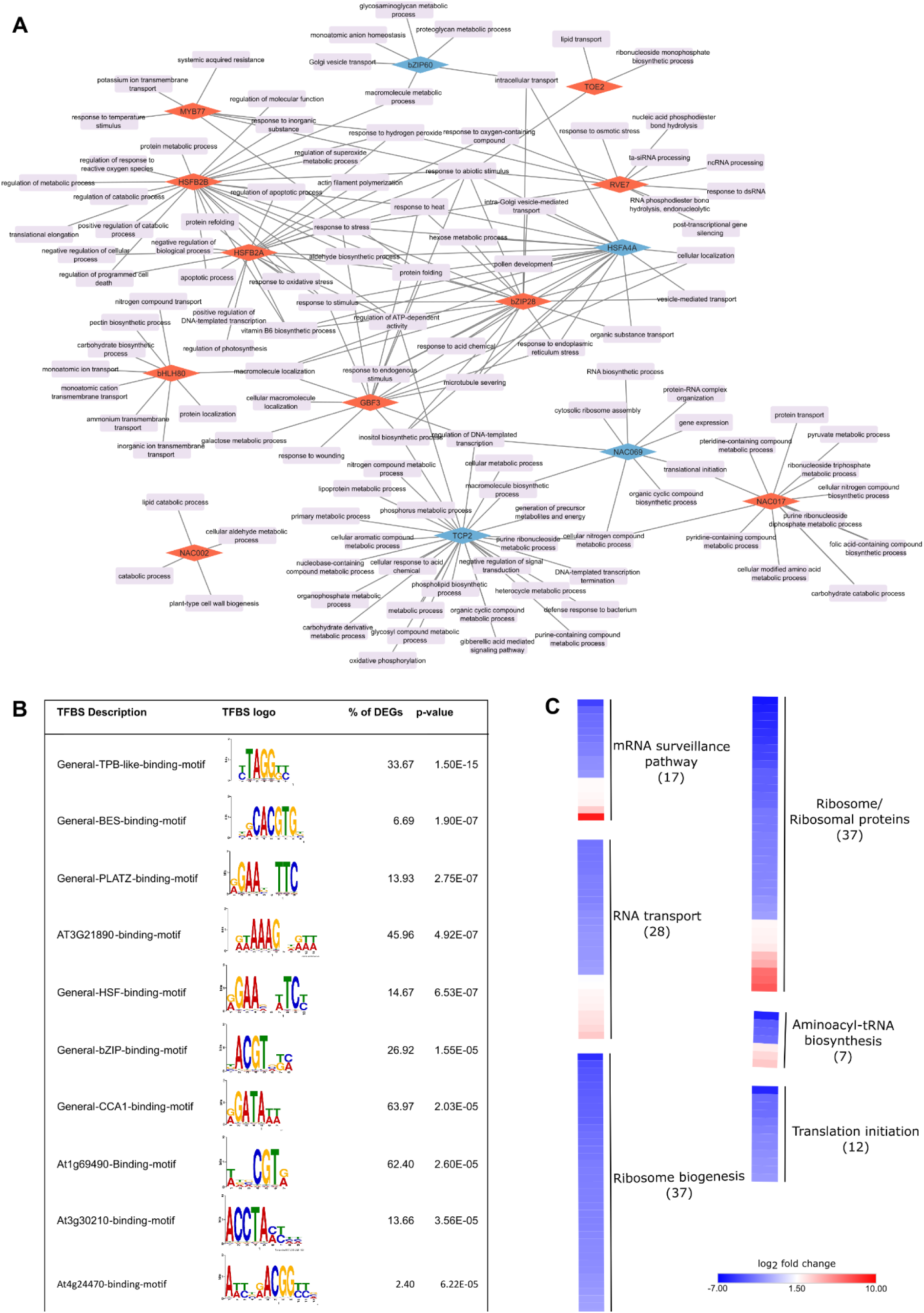
**(A)** Enriched TFs and biological processes regulatory networks in heat stressed microspores. Fourteen transcription factors (diamond boxes) were identified as significantly enriched and their targets genes overrepresented in the differentially expressed gene sets in microspores in response to heat stress in microspores. Red diamond box represents up-regulated TF, and blue diamond box represents downregulated TF. **(B)** Details of TFBS enrichment analysis in the promoters of differentially expressed genes in heat-stressed microspore. **(C)** Heatmap representation of log_2_ foldchange in expression of heat stress responsive genes associated with post transcriptional (mRNA surveillance pathway, mRNA transport) and translational machinery (Ribosome biogenesis, Ribosomal proteins, aminoacyl; t-RNA biosynthesis and translation initiation) in microspores.

The homolog of bZIP28 was upregulated by 1.4 log_2_ fold, whereas bZIP60 was downregulated by -2.3 log_2_ fold in heat-stressed microspores. This opposite regulation of bZIP28 and bZIP60 contrasts with the upregulation of both bZIP TFs in reproductive tissues in Arabidopsis (S.-S. Zhang et al., 2017). Also in Arabidopsis, Poidevin et al. (2021), reported no significant change in expression of bZIP60 in germinated pollen in response to heat stress. The upregulation of bZIP60 in response to heat stress is also reported in leaf lamina in maize, establishing role of bZIP60 in linking the unfolded protein response (UPR) and heat stress response (Li et al., 2020). The observed differences across heat stress transcriptome studies and the regulation of bZIP60 and bZIP28 are probably due to differences in tissue analysed. Furthermore, the similarity between *AtbZIP28* gene and its homolog in *B. napus* (*BnaA09g24670D*) is 67.9% and that of *AtbZIP60* and its *B. napus* homolog (*BnaC08g05600D*) is 55%, indicating probable neo-functionalization over the course of evolution. Another enriched bZIP TF, GBF3 (*BnaA05g01520D*), in heat-stressed microspores indicates the involvement of abscisic acid (ABA) mediated signalling (Dixit et al., 2019; Ramegowda et al., 2017) in heat stress response during pollen development. However, the role of GBF3 in male reproductive development in plants has not yet been reported.

Next, to identify enriched transcription factor binding sites (TFBS), we performed promoter motif analysis using the 1kb promoter sequences of the heat responsive genes in microspores (**Figure 3B**). ‘General-HSF-binding-motif’ and ‘General-PLATZ-binding-motif’ which act as heat stress elements, were present in promoters of ∼14% of the heat responsive genes. HSFs are well known to bind to the heat stress elements (HSEs: 50-nGAAnnTTCnnGAAn-30 or 50-nTTCnnGAAnnTTCn-30) present in the promoters of their target genes (Guo et al., 2016). A considerably higher percentage of differentially expressed genes promoters had TFBS recognised by MYB (‘General-TPB-like-binding-motif’, ‘General-CCA1-binding motif’, ‘At3g30210 - binding-motif’) and NAC (‘At1g69490 -binding-motif’) TFs.

In addition to transcriptional changes, pre-mRNA alternative splicing (AS) is another important mechanism involved in heat stress response (Laloum et al., 2018; Ling et al., 2021). In the present study, a total of 58 differentially expressed genes were associated with the term “Spliceosome” out of which 36 were upregulated and 21 were downregulated (**Table S4**). Genes homologous to SMP1 which encodes CCHC zinc finger proteins with similarities to step II splicing factors involved in 3’ splice site selection were significantly upregulated in heat-stressed microspores. We also identified differential regulation of DEAD-box RNA helicases (RH); for example, RH42 homologs were upregulated in heat-stressed microspores. Numerous studies have highlighted the role of DEAD-box RNA helicases in enhancing tolerance to abiotic stress in transgenic plants (Nidumukkala et al., 2019; Tuteja et al., 2014; Yarra & Xue, 2020). Recently, a DEAD-box RNA helicase located in the nucleolus, identified as THERMOTOLERANT GROWTH REQUIRED 1 (TOGR1), was discovered through map-based cloning in rice (Wang et al., 2016). TOGR1 has been shown to contribute to heat tolerance by interacting with the pre-rRNA processing complex and stabilizing ribosomal RNA levels under high-temperature conditions by increasing helicase activity (Wang et al., 2016). Additionally, the expression of OsTOGR1 in Chinese cabbage resulted in improved thermotolerance and the upregulation of several heat-responsive genes (Yarra & Xue, 2020). This might suggest a similar involvement of DEAD-box RNA helicases in heat tolerance mechanisms within Brassica species.

Other key genes associated with heat stress induced mis-regulation of the mRNA processing machinery are the homologs of SERINE/ARGININE-RICH PROTEIN SPLICING FACTOR 30 (SR30) protein encoding genes. SR30 homologs were significantly upregulated in heat-stressed microspores. The diversity and presence of splicing factors, such as SR proteins, is potentially crucial for determining the splicing patterns of heat response genes, including HSFs and HSPs as they are the core genes involved in alternative splicing (Rosenkranz et al., 2022) and in mammals SR proteins are involved in repression of alternative splicing in response to heat stress (Shin et al., 2004). Furthermore, differentially expressed HSP70 proteins potentially impact mRNA metabolism through multiple mechanisms, such as aiding in the folding of emerging polypeptide chains, activating signalling pathways, facilitating the resolution of stress granules, and selectively degrading mRNA (Maruri-López et al., 2021; Walters & Parker, 2015; T.-Y. Wang et al., 2021). This extensive suppression of pre-mRNA splicing is not only attributed heat stress induced damages but represents a strategic regulatory adaptation that aids organisms in surviving heat stress. Taken together, these findings provide a detailed understanding of the transcriptional and post-transcriptional mechanisms activated in microspores under heat stress.

### Coordination of Protein Homeostasis and Translational Control is Altered in Heat-Stressed Microspores

Controlled and coordinated translation of mRNA is vital for plant developmental processes including gametogenesis (Huang et al., 2021). In our study, we identified more than hundred genes associated with the functional pathway category “Translation” (**Figure 2F**). This category includes genes involved in the “Aminoacyl-tRNA biosynthesis”, “mRNA surveillance pathway”, “Ribosome”, “Ribosome biogenesis in eukaryotes” and “RNA transport” pathways (**Figure 3C**) with majority of these genes significantly downregulated in heat-stressed microspores (**Figure 2D**, **3C**). The identified differentially expressed genes were homologous to several eukaryotic translation initiation factors (eIF), Nucleic acid binding, OB fold like proteins, genes encoding ribonucleoprotein complex subunits, ribosomal proteins, pre-RNA processing ribonucleoproteins and several other genes mediating different aspects of the translational machinery (**Table S5**). Interestingly, in Arabidopsis, young leaves exposed to 1h of heat stress had significantly increased transcripts for genes related to, ribosome biogenesis, and mRNA surveillance, highlighting differences in heat stress response between reproductive and vegetative cells/tissues (Xiang & Rathinasabapathi, 2022).

Alterations in the regulation of genes linked to the mRNA surveillance pathway can impact its primary function, which is to prevent the translation of faulty mRNA that could lead to the production of harmful proteins (Behm-Ansmant et al., 2007). In heat-stressed microspores, several Serine/threonine-protein phosphatases, including homologs of TYPE ONE PROTEIN PHOSPHATASES (TOPPs), were differentially expressed in *B. napus*. TOPPs encoding genes known to influence various biological processes, abscisic acid (ABA) signalling (Zhang et al., 2020), autophagy (Wang et al., 2022) and maintaining cell wall integrity during the tip-growth of pollen tubes and root hairs (Franck et al., 2018). Additionally, homologs of the TOPP genes in rice (*Oryza sativa*), wheat (*Triticum aestivum*), and soybean (*Glycine max*) are reported to be involved response to abiotic stresses, such as salt stress in rice and wheat, and drought in soybean (Bradai et al., 2018; Liao et al., 2016; S. Wang et al., 2021).

Another key gene involved in the mRNA surveillance pathway that was downregulated in heat-stressed microspores is POLYADENYLATE-BINDING PROTEIN 5 (PAB5). Its ortholog in Arabidopsis is also downregulated in the *ams* mutant, which are characterized by an expanded tapetal layer and aborted microspores (Sorensen et al., 2003). This phenotypic similarity suggests that the downregulation of PAB5 in *B. napus* in response to heat stress may contribute to microspore abortion, possibly through a mechanism comparable to that observed in Arabidopsis.

Among the significant DEGs, thirty-eight genes related to ribosome biogenesis, along with twelve translation initiation factors and subunits, were found to be significantly downregulated in heat-stressed microspores (**Figure 3C**, **Table S5**). This suggests that heat stress in uninucleate microspores leads to a notable inhibition of translation. Ribosome biogenesis factors (RBFs) are known to play essential roles in reproductive development (Missbach et al., 2013), and certain translation initiation factors, such as eIF5B, have been shown to be critical for heat stress tolerance in plants (Son & Park, 2023; L. Zhang et al., 2017). Heat stress-induced inhibition of translation can also prompt the formation of stress granules (SGs), membraneless aggregates composed of untranslated mRNAs, translation initiation factors, RNA-binding proteins, and other cellular components (Chantarachot & Bailey-Serres, 2018; Kosmacz et al., 2019; Tong et al., 2022; Yan et al., 2022). SGs play a crucial role in regulating the stress response by selectively storing mRNAs and proteins, thereby protecting them during periods of stress (Merret et al., 2013; Merret et al., 2015; Takahara & Maeda, 2012). This sequestration mechanism effectively pauses translation and protein activity, with SG disassembly allowing for a rapid return to normal cellular functions once the stress subsides. The observed downregulation of translation in heat-stressed microspores may also represent a cellular strategy to cope with the adverse effects of heat stress.

Exposure to temperatures exceeding optimal growth conditions leads to misfolding and aggregation of a wide range of proteins. Consequently, cells have developed robust, compartmentalized stress response mechanisms known as the “Unfolded Protein Response” which adapts to the extent of protein misfolding, ensuring the maintenance of protein homeostasis. “Protein processing in ER” was the most enriched KEGG pathway and “protein folding” was the most significantly upregulated GO term in heat-stressed microspores. Significant enrichment of GO terms related to response to heat, protein folding, and ER unfolded protein response demonstrated the ability of microspores to sense and respond to high temperature in a similar manner to ER-dependent stress responses in flower and vegetative tissues in plants (S.-S. Zhang et al., 2017). We mined the literature to identify the various gene components of UPR in ER (Singh et al., 2021) and then identified their homologous genes in *B. napus* (**Table S6**). In Arabidopsis, bZIP28 and bZIP60 mediated UPR is crucial for heat responses in reproductive tissues (S.-S. Zhang et al., 2017). As discussed earlier, in heat-stressed microspores these bZIP TFs displayed antagonistic regulation and differentially regulated the gene expression of a plethora of target genes associated with heat stress and development related biological processes (**Figure 3A**).

Furthermore, an array of proteins that act as molecular chaperones or foldases mediate protein folding in ER. Genes homologous to molecular chaperones such as BiP1, SHD, HSP20, HSP21 and several small HSPs were significantly upregulated. Few HSP homologs, for example BiP3, HSP14.7, HSP20-like chaperone and HSP70 were downregulated in heat-stressed microspores. Heat stress exposure also significantly upregulated homologs of calcium binding chaperone (CNX1, CRT1, CRT2 and CRT3) in *B. napus* microspores. Foldases such as protein disulfide isomerases (PDIs) catalyse the formation and isomerization of disulfide bonds to assist the folding of misfolded proteins. PDI9, PDI10 and PDI11 homologs were also upregulated in heat-stressed microspores. As discussed earlier PDI proteins play an integral role in ER stress, UPR and are also associated with male reproductive development and heat stress response (Feldeverd et al., 2020; Q. Zhao, X. Guan, L. Zhou, Y. Xu, et al., 2023), add rice reference, Furthermore, in heat-stressed microspores, genes homologous to the DnaJ proteins (ERDJ3A, TMS1, ERDJ3B, ERDJ2B) were significantly upregulated. TMS1 (*Thermosensitive Male Sterile 1*) is reported to play a significant role in regulating pollen thermotolerance (Ma et al., 2015) and defective interaction of AtERdj3B with BiP results in heat stress-sensitive fertility defect in Arabidopsis (Yamamoto et al., 2020).

Autophagy is an essential pathway for quality control of the ER, facilitating the removal of damaged or unnecessary materials and the recycling of cellular components under developmental or environmental stress. In heat-stressed microspores, homologs of various genes encoding autophagy-related proteins (ATGs) were identified, with elevated temperatures causing a decrease in the expression of ATG1b, ATG6, ATG8a, ATG8e, ATG8f, ATG8h, ATG13b, ATG18e, ATG18f, ATG101, and an increase in ATG18a and ATG18h (**Table S6**). The upregulation of ATG18a in heat stressed microspore is a consequence of heat induced ER stress which triggers ER-phagy and the transport of the ER to the vacuole for degradation is dependent on the ATG18a protein (Liu et al., 2012). Additionally, the differential regulation of ATGs in response to diverse abiotic stresses is reported in *B. napus* and several other plant species (Eshkiki et al., 2020). Among other ATGs, ATG8 is a component of the ATG conjugation systems, which attach to the membranes of autophagosomes to aid in the formation of autophagosomes and the selective degradation of cargo (Bu et al., 2020). Recently, Zhou et al. (2023) reported unexpected function of Arabidopsis ATG8 together with its interactor CLC2 (CLATHRIN LIGHT CHAIN 2) in the reestablishment of the Golgi apparatus following heat shock, independent of the autophagic precursors. Interestingly the repression of ATG8 homologs and upregulation of the CLC2 homolog in heat stressed *B. napus* microspores suggests a disruption in the Golgi’s restoration process. However, only few reports have highlighted the role of autophagy in plant male reproductive development. ATG5 and ATG7-dependent autophagy contributes to the removal of cytoplasm during the differentiation of sperm cells in lower plants, producing motile sperm for fertilization (Sanchez-Vera et al., 2017). Similarly, in rice, autophagy is vital for the development of the tapetum; ATG7 mutants exhibited male sterility due to the buildup of lipid bodies and organelles (Kurusu et al., 2014). Furthermore, in heat stressed microspores, BAG6 (an autophagy related gene) was significantly upregulated, which potentially impacts male reproductive success, as demonstrated by the reduced seed production and severe pollen defects in BAG6 overexpressing *Fragaria vesca* (F. Zhao et al., 2023).

Proteins are exported from ER to the Golgi apparatus for sorting to their final destination. Based on functional annotation analysis, GO terms such as “intra-Golgi vesicle mediated transport”, “vesicle-mediated transport” and “intracellular protein transport” along with KEGG pathways “protein export” and “SNARE interactions in vesicular transport” were identified, which indicate the high temperature mediated misregulation of protein transport from the ER to the Golgi apparatus in microspores (**Table S6**). Detailed understanding of the intersection of functionally related yet-distinct pathways that connect with protein transport during pollen development and heat stress response is required. The differential regulation of genes involved in protein folding, sorting, degradation, and transport is not just the consequence of heat stress response but also plays critical role in progression of pollen development (Singh et al., 2021).

### Gene Involved in Cell Cycle Control and Progression of Pollen Development are Mis-regulated in Heat-Stressed Microspores

The completion of male meiosis marks the initiation of a unique pathway of cellular differentiation that leads through a simple cell lineage to the formation of the male gametophytes – pollen grains – in flowering plants. Two sequential but different modes of mitotic divisions, pattern the male gametophyte. In Arabidopsis, alterations in cell cycle regulators can disrupt the progression of mitosis, leading to gametophytic fatality (Takatsuka et al., 2015). This prompted us to perform literature mining to identify genes that participate in asymmetrical division and cell mitosis, followed by identification of their homologs in heat stressed responsive genes in *B. napus* microspores (**Figure 4A**, **Table S7A**).

**Figure 4.**
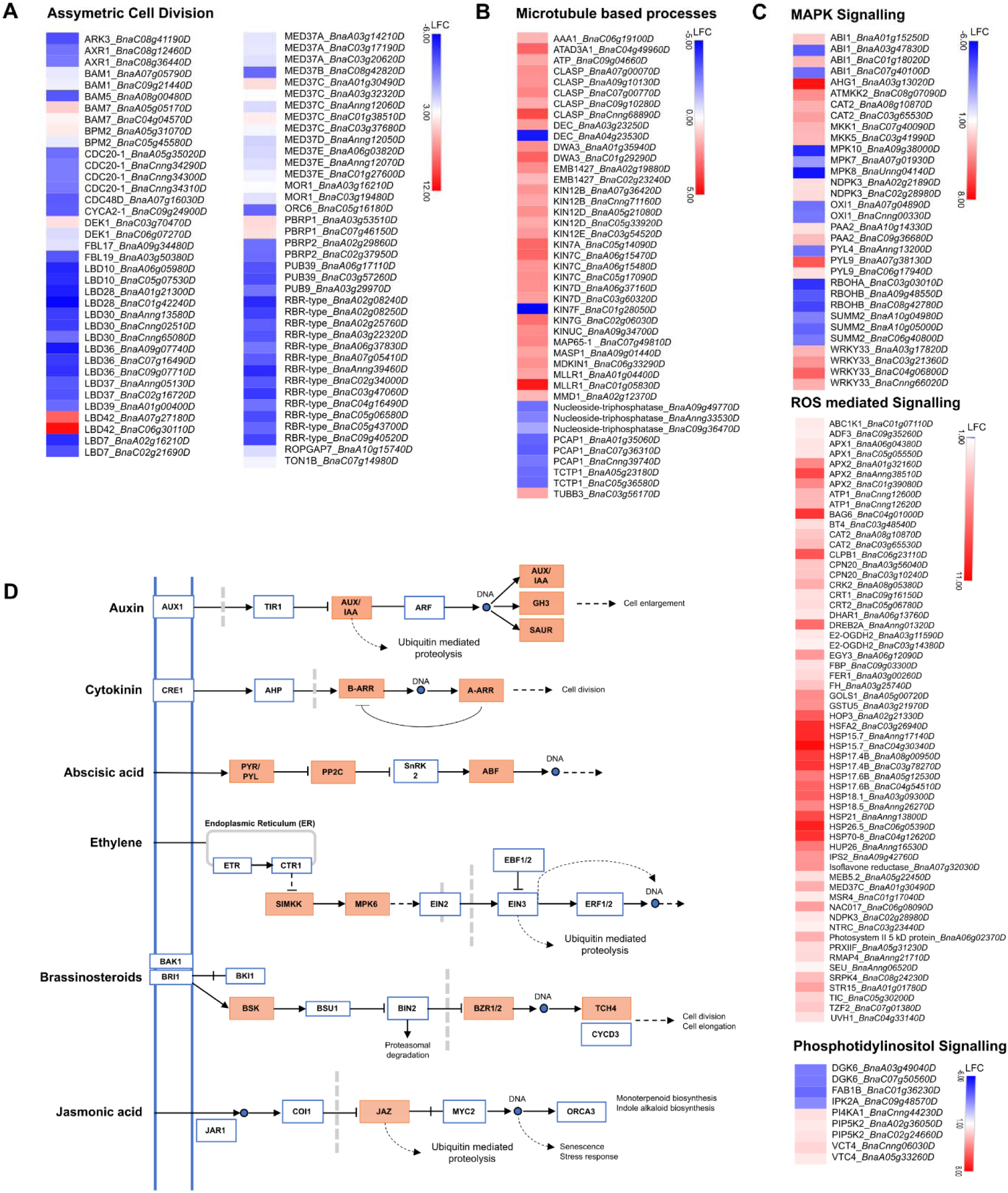
Heatmap representation of log2 foldchange in expression of heat stress responsive genes associated with asymmetric cell division **(A)**, microtubule-based processes **(B)**, and signalling pathway namely MAPK signalling, ROS-mediated signalling, and Phosphatidylinositol signalling **(C)**. **(D)** Differentially expressed gene-enrichment of the plant hormone signal transduction pathway by KEGG annotation. The key regulatory components in multiple hormone response pathways are presented as their names (solid brown coloured boxes represent differentially regulated genes in heat-stressed microspores).

SIDECAR POLLEN/LATERAL ORGAN BOUNDARY 27 (SCP/LBD27) and LBD10 are required for pollen development and for maintaining division asymmetry during PMI (Kim et al., 2015). In heat-stressed microspores, probable LBD10 homologs are downregulated. However, we did not observe differential regulation of SCP/LBD27 homologous genes in *B. napus* heat-stressed microspores. *B. napus* genes homologous to CELL DIVISION CYCLE 20.1 (CDC20.1) and CDC48D were significantly downregulated in heat-stressed microspores. CDC20.1 encodes a protein that interacts with APC subunits, components of the mitochondrial checkpoint complex and mitotic cyclin substrates and is indispensable for normal plant development and fertility (Niu et al., 2015). CDC48 is reported to be critical for cytokinesis, cell expansion and differentiation in plants (Rancour et al., 2002) and it also plays essential roles in ERAD pathway (Christianson et al., 2023). Another important cell cycle relating gene, TON1B was upregulated in heat-stressed microspores. TONNEAU 1B (TON1B) plays a role in the organization of microtubule arrays at the centrosome through interaction with CENTRIN1 (CEN1) (Drevensek et al., 2012).

Furthermore, asymmetrical cell division relies on microtubules, and the majority of identified mutants impacting this process are associated with microtubule function (Twell, 2011). In heat-stressed microspores differentially expressed genes were also associated with “microtubule-based processes” and “regulation of microtubule cytoskeletal organisation” (**Figure 4B**, **Table S7B**). For instance, CLIP-ASSOCIATED PROTEIN (CLASP) encoding genes in *B. napus* were upregulated in heat-stressed microspores. CLASP is a microtubule-associated protein that is involved in both cell division and cell expansion (Ambrose et al., 2007). TUBB3 gene encodes tubulin beta-3 chain protein which is a major constituent of microtubules (Ben-Nissan et al., 2008); its ortholog in *B. napus* microspores is downregulated in response to heat stress. MALE MEIOCYTE DEATH 1 (MMD1), which is required for microtubule organization and regulation of cell cycle transitions (Andreuzza et al., 2015) was upregulated in heat-stressed microspores. These results support our microscopic observations and provide the details of transcriptional reprogramming underlying the failure of heat stressed microspores in undergoing PMI and PMII, and, resulting in aberrant pollen development in *B. napus*.

### Heat stress alters multiple signalling pathways critical for pollen development

Crosstalk between multiple signal transduction and signalling pathways tightly regulate the progression of male reproductive development. One such pathway is the MAPK signalling pathway, which is involved in male reproductive development and other aspects of plant reproductive processes (Komis et al., 2018). MAPK signalling cascade integrates and channels signal transduction for the expression of stress-responsive genes mediated through phosphorylation. In heat-stressed microspores, orthologs of MAP KINASE KINASE 2 (MKK2), MKK1, MKK5 were upregulated whereas MPK7, MPK8, MPK10 were downregulated highlighting the misregulation of MAPK cascades in response to high temperature during male reproductive development (**Figure 4C**, **Table S8A**). WRKY33, a target of MAPK signalling was also upregulated in heat-stressed microspores. In tomato, similar elevation of WRKY33 transcripts were reported in response to 4h of heat stress exposure at 45 (Zhou et al., 2014). On the contrary, repressed expression of WRKY33 is reported in Arabidopsis seedlings in response to 1-2h of heat stress treatment at 42-48 (Li et al., 2011) highlighting tissue and species-specific response to acute heat stress. Furthermore, orthologs of SUPPRESSOR OF MKK1 MKK2 2 (SUMM2) are significantly downregulated in heat-stressed microspores. SUMM2 is associated with the regulation of cell death during sexual reproduction in plants, and it interacts with the MKK4 which is phosphorylated by MEKK1 and MKK1/MKK2 proteins (Völz et al., 2022).

Under high temperature the balance between production and detoxification is disturbed, resulting in the build-up of ROS (Devireddy et al., 2021). The accumulation of ROS is considered as one of the earlier responses in the heat stress sensing pathways. In heat-stressed microspores, we identified more than sixty genes associated with “response to oxidative stress” all of which were significantly upregulated except for DEHYDROASCORBATE REDUCTASE 3 (DHAR3), RESPIRATORY BURST OXIDASE HOMOLOGUE A (RBOHA) and RBOHB. Upregulated *B. napus* orthologs of key ROS scavenging enzymes such as ASCORBATE PEROXIDASE 1 (APX1/MEE6), APX2, CATALASE (CAT2), DHAR1 and GOLS1 highlighted the heat triggered response of attempting to maintain ROS homeostasis in microspores (**Figure 4C**, **Table S8B**). Studies have indicated that the indirect activation of MAPK signalling through ROS mediation triggers a feedback loop, resulting in a reduction of cellular ROS levels as a consequence of MAPK signalling. In Arabidopsis, regulation of CAT2 expression is dependent on activation of MAPK cascade. In addition to modulating the expression of genes involved in ROS scavenging, MAPK signalling is also associated with the suppression of RBOH expression. Signalling of ROS independent of MAPK is initiated through the activation of redox sensor proteins, leading to the subsequent activation of enzymes responsible for scavenging ROS. Additionally, key TFs such has NAC017, which is upregulated in heat stressed microspores, is reported to be released from the ER in response to heat triggered elevated levels of hydrogen peroxide H_2_O_2_ (Ng et al., 2013).

The ROS and MAPK signalling coupled flexible feedback loop also serves to amplify the effects of hormonal signalling in response to stress in plants (Devireddy et al., 2021). In heat-stressed microspores, several genes associated with phytohormone-mediated signalling were identified to be involved in auxin, cytokinin, brassinosteroids, jasmonic acid (JA), ABA and ethylene signalling (**Figure 4D**, **Table S8C**). Key component and repressor of the ABA signalling pathway, ABA INSENSITIVE (ABI) ortholog genes were differentially regulated in heat-stressed microspores. ABA-HYPERSENSITIVE GERMINATION 1 (AHG1) gene which acts as a negative regulator of ABA responses, was also significantly upregulated in heat-stressed microspores. Additionally, in auxin signalling, INDOLE-3-ACETIC ACID INDUCIBLE 9 (IAA9) homologs were upregulated, SMALL AUXIN UPREGULATED RNA 70 (SAUR70) was downregulated and auxin responsive GH3 family protein homologs were differentially regulated. Contrary to vegetative tissues, heat stress has been shown to result in a decline in auxin content in anthers of several crops (Min et al., 2014; Sakata et al., 2010). Interestingly in barley, when applied exogenously, auxin treatment has been shown to enhance pollen thermotolerance during early development stages in response to moderate heat stress (Sakata et al., 2010).

Furthermore, two key genes associated with brassinosteroid (BR) signalling were differentially regulated in heat-stressed microspores (**Table S8C**). BEH1 was upregulated and BES1 were downregulated in heat-stressed microspores. Two BSK2 homologs which acts as positive regulator of BR signalling downstream of the receptor kinase BRI1, were significantly downregulated in heat-stressed microspores. These results indicate possible negative regulation of BR accumulation and signalling in heat-stressed microspores. BRs control male fertility at least in part via directly regulating key genes (SPL/NZZ, TDF1, MS2, MYB103, MS1, and AT3G23770) for anther and pollen development in Arabidopsis (Ye et al., 2010). Recently, the role of BZR1-dependent ROS production playing a critical role in the BR-mediated regulation of tapetal cell degeneration and pollen development in tomato plants was also reported (Yan et al., 2020).

Phosphoinositides are a class of cellular signalling lipids that are derived from phosphatidylinositol (PtdIns) by the action of lipid kinases and phosphatases and are reported to activate within minutes of a sudden temperature rise in Arabidopsis (Mishkind et al., 2009). In sorghum, changes in pollen phospholipids and an increase in ROS levels in response to high night temperatures is potentially related to the reduction in pollen function (Prasad & Djanaguiraman, 2011). In heat-stressed microspores in *B. napus*, we identified nine differentially expressed genes associated with the PtdIns signalling system (**Figure 4C**, **Table S8D**). Genes encoding DIACYLGLYCEROL KINASE 6 (DGK6), were downregulated in heat-stressed microspores. DGKs convert DAG to Phosphatidic acid (PA) and also act as putative signalling component between PA signalling and Ca^2+^ signalling (Kue Foka et al., 2020). FAB1B homologs were also downregulated in heat-stressed microspores. FAB1B act as a 1-phosphatidylinositol-3-phosphate (PtdIns3P) 5-kinase (Koldenkova & Hatsugai, 2017). FAB1B and FABID show highest level of expression in Arabidopsis pollen when compared to the rest of the FAB gene family members. These genes are suggested to play a complementary role in the regulation of membrane recycling, vacuolar pH and homeostatic control of ROS in pollen tube growth (Serrazina et al., 2014). Downregulation of FAB1B in heat-stressed microspores might be involved in negative regulation of pollen development.

Taken together, these results highlight that heat stress alters the intricate and complex network of multiple signalling pathways potentially leading to disruption of regulatory pathways underlying pollen development.

## Conclusion

This study elucidates the profound impact of short-term heat stress on *B. napus* pollen development, particularly during the critical uninucleate microspore stage. Acute heat exposure disrupts the normal progression of pollen development, leading to significant fertility reduction through failed mitotic divisions and subsequent pollen abortion. Employing high-throughput RNA sequencing, we identified the heat-responsive transcriptional reprogramming and genes, highlighting the activation of classical heat stress responses and the differential regulation of novel genes potentially critical for heat stress resilience. Key pathways such as protein folding, sorting, degradation, and genetic information processing were notably affected, along with the regulation of heat shock factors and proteins, phytohormones, ROS scavengers, metabolic genes, phosphatidylinositol signalling, cell cycle, and translational machinery. The evident crosstalk among these pathways underscores the complexity of the heat stress response in pollen development. Our findings lay the groundwork for future studies aimed at enhancing the thermotolerance of *B. napus*, offering insights into potential genetic targets for improving crop resilience to climate-induced thermal stress.

## Data availability statement

The original contributions presented in the study are included in the article/Supplementary Material. The RNA-Seq datasets presented in this study are deposited at the NCBI Sequence Read Archive (BioProject ID: PRJNA666230). Further inquiries can be directed to the corresponding author.

## Author contributions

NL conducted the experiments, analysed the sequencing data, and prepared a draft of the manuscript. PB and MS conceived the research, supervised, and extensively edited the article. All authors contributed to the article and approved the submitted version.

## Acknowledgments

We acknowledge the University of Melbourne for supporting NL through a Melbourne Postgraduate Research Scholarship.

## Supplementary Files

**Supplementary Figure 1. (A)** Schematic representation of the temperature regime followed for exposing the plants to a single episode of heat stress (40°C for 4h); das: days after sowing. **(B)** PCA plot to illustrate the biological relatedness of replicates for each sample (NS: non-stressed, HS: heat-stressed). **(C)** Venn diagram representing the intersecting and specific differential regulation of heat responsive transcriptome between microspore and mature pollen.

**Supplementary Figure 2.** Heatmap representation of expression of differentially expressed genes identified as transcription factors (TF) belonging to different TF families. Insets of four important heat stress responsive TF families are provided to highlight the annotated members of these TF family that regulate the transcriptional cascade in heat-stressed microspores. NS_Mic: Non-stressed microspores, HS_Mic: Heat-stressed microspores.

**Table S1.** Results table generated by DESeq2 analysis for “heat-stressed microspores vs non-stressed microspores” summarizing the base means across samples, log_2_ fold changes, standard errors, and test statistics for genes which pass an adjusted p value threshold of 0.01.

**Table S2.** Number of differentially expressed genes in heat-stressed microspores annotated using multiple approaches.

**Table S3.** Summary of transcription factor (TF) prediction and enrichment analysis. **(A)** Details of all TFs identified as heat responsive in microspores, **(B)** Details of enriched TFs and the number of their differentially expressed target genes.

**Table S4.** Details of the heat stress responsive gene in microspores associated with “Spliceosome” pathway.

**Table S5.** Details of the heat stress responsive gene in microspores associated with translation related pathways.

**Table S6.** Details of the heat stress responsive gene in microspores associated with protein homeostasis pathways: UPR: Unfolded Protein Response, Autophagy, and Protein Transport.

**Table S7.** Details of the heat stress responsive gene in microspores associated with **(A)** asymmetric cell division and **(B)** microtubule related processes.

**Table S8.** Details of the heat-stressed microspore DEGs associated with **(A)** MAPK signaling cascade, **(B)** Phytohormone mediated signaling, **(C)** ROS mediated signaling, and **(D)** Phosphatidylinositol signaling pathways”

