## Supplementary Figures for "Heat-Triggered Transcriptional Reprogramming in Microspores Disrupts Progression of Pollen Development in *Brassica napus* L"

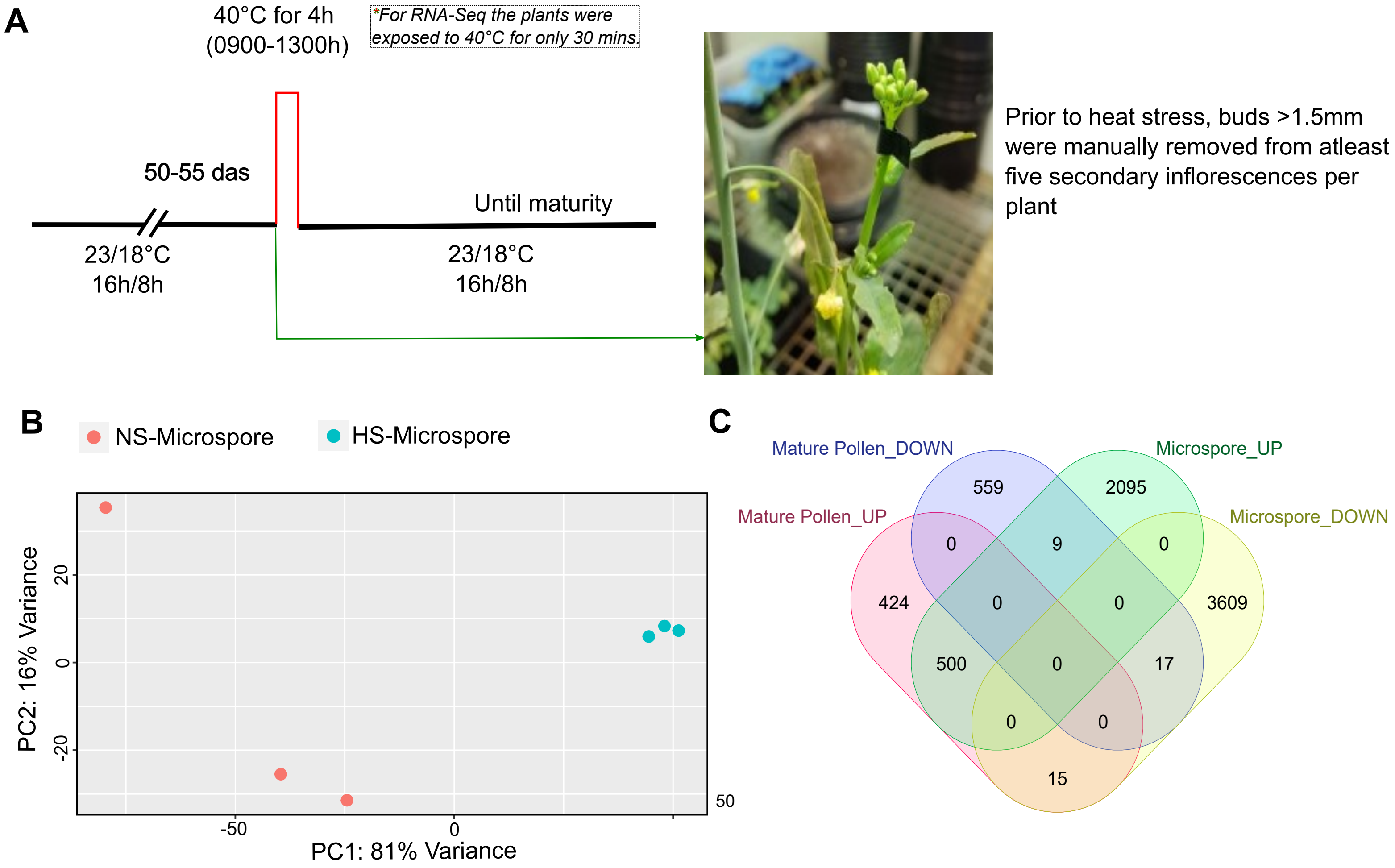


**Supplementary Figure 1. (A)** Schematic representation of the temperature regime followed for exposing the plants to a single episode of heat stress (40°C for 4h); das: days after sowing. **(B)** PCA plot to illustrate the biological relatedness of replicates for each sample (NS: non-stressed, HS: heat-stressed).**(C)** Venn diagram representing the intersecting and specific differential regulation of heat responsive transcriptome between microspore and mature pollen.

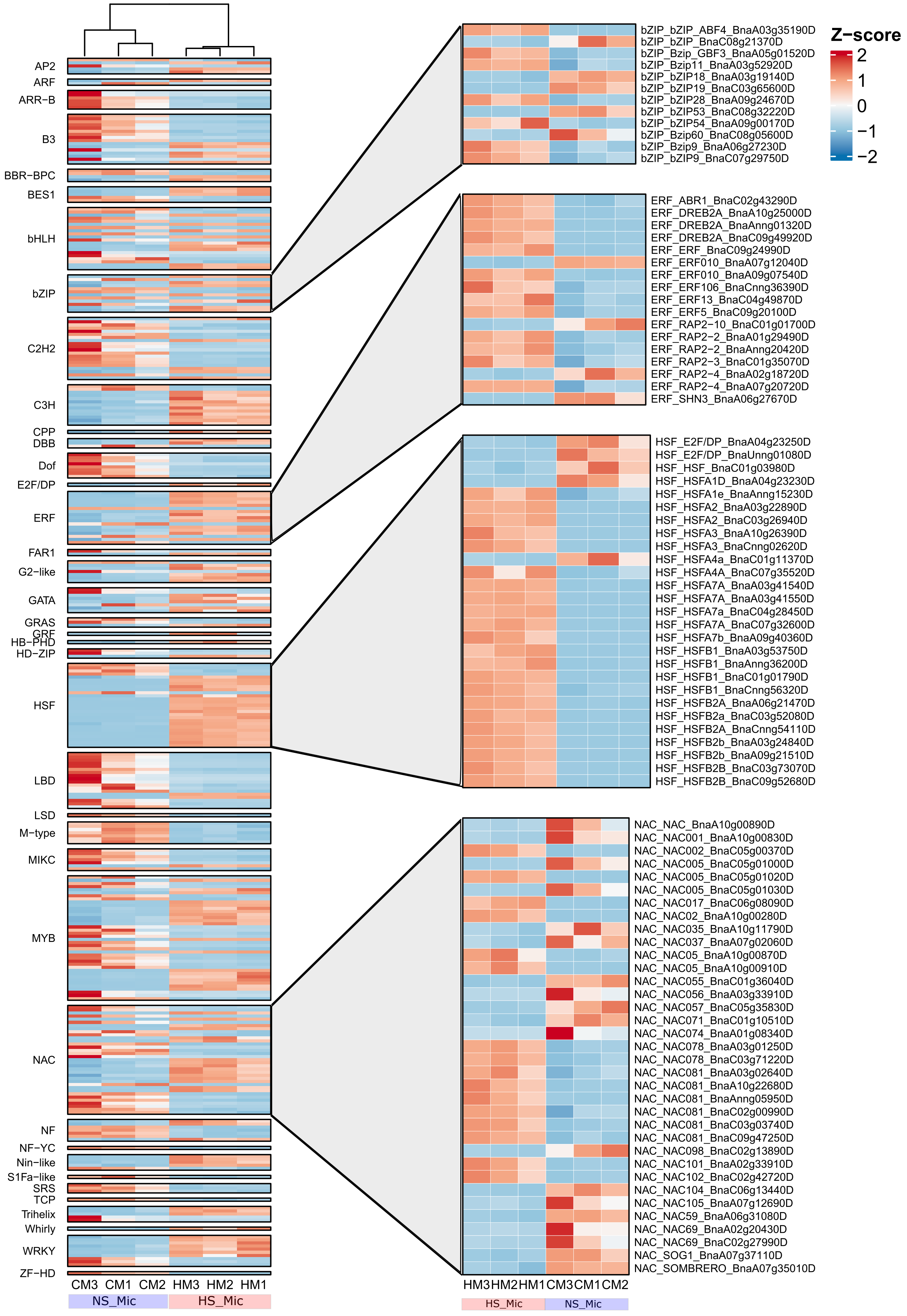


**Supplementary Figure 2.** Heatmap representation of expression of differentially expressed genes identified as transcription factors belonging to different TF families. Insets of four important heat stress responsive TF families is provided to highlight the annotated members of these TF family that regulate the transcriptional cascade in heat-stressed microspores. NS_Mic: Non-stressed microspores, HS_Mic: Heat-stressed microspores.
